# Are all global alignment algorithms and implementations correct?

**DOI:** 10.1101/031500

**Authors:** Tomáš Flouri, Kassian Kobert, Torbjørn Rognes, Alexandros Stamatakis

## Abstract

Pairwise sequence alignment is perhaps the most fundamental bioinformatics operation. An optimal global alignment algorithm was described in 1970 by Needleman and Wunsch. In 1982 Gotoh presented an improved algorithm with lower time complexity. Gotoh’s algorithm is frequently cited (1447 citations, Google Scholar, May 2015), taught and, most importantly, used as well as implemented. While implementing the algorithm, we discovered two mathematical mistakes in Gotoh’s paper that induce sub-optimal sequence alignments. First, there are minor indexing mistakes in the dynamic programming algorithm which become apparent immediately when implementing the procedure. Hence, we report on these for the sake of completeness. Second, there is a more profound problem with the dynamic programming matrix initialization. This initialization issue can easily be missed and find its way into actual implementations. This error is also present in standard text books. Namely, the widely used books by Gusfield and Waterman. To obtain an initial estimate of the extent to which this error has been propagated, we scrutinized freely available undergraduate lecture slides. We found that 8 out of 31 lecture slides contained the mistake, while 16 out of 31 simply omit parts of the initialization, thus giving an incomplete description of the algorithm. Finally, by inspecting ten source codes and running respective tests, we found that five implementations were incorrect. Note that, not all bugs we identified are due to the mistake in Gotoh’s paper. Three implementations rely on additional constraints that limit generality. Thus, only two out of ten yield correct results. We show that the error introduced by Gotoh is straightforward to resolve and provide a correct open-source reference implementation. We do believe though, that raising the awareness about these errors is critical, since the impact of incorrect pairwise sequence alignments that typically represent one of the very first stages in any bioinformatics data analysis pipeline can have a detrimental impact on downstream analyses such as multiple sequence alignment, orthology assignment, phylogenetic analyses, divergence time estimates, etc.

## 1 Introduction

The *Needleman-Wunsch* (NW) [12] and *Smith-Waterman* [18] algorithms for computing optimal global and local alignments are among the most important algorithms in bioinformatics and computational biology. They are typically presented in undergraduate lectures at many computer science and bioinformatics departments around the globe. Although Needleman and Wunsch described their algorithm in their seminal paper in 1970, the algorithm had already been discovered several times before. In fact, Damerau and Levenshtein independently described the algorithm in 1964 [4] and 1965 [10]. Analogous algorithms with quadratic run-times were also independently developed by Vintsyuk in 1968 for speech processing [20], and in 1974 by Wagner and Fischer for string matching [21]. In 1972, Sankoff presented an improved dynamic programming algorithm with quadratic time complexity for this problem by making additional assumptions [15]. The algorithm by Sankoff maximizes the number of matches between two sequences, without penalizing gaps. Needleman and Wunsch described their algorithm in terms of maximizing similarity between two sequences. Levenshtein described the problem in terms of minimizing the *edit distance*, that is, the cost of edit operations (insertion, deletion, substitution) for transforming one sequence into another. In 1974, Sellers showed that these two variations are in fact equivalent [16]. Finally, in 1982 Gotoh presented a quadratic time algorithm to compute global sequence alignments with affine gap penalties [8]. Note that, Gotoh’s approach also reduces the time complexity of the Smith-Waterman *local* alignment algorithm. While the underlying idea of Gotoh’s algorithm is valid and can yield the optimal pairwise sequence alignment, there are two issues that can lead to erroneous, that is, sub-optimal, alignments based on Gotoh’s original description. The first issue (*index issue*) is straight-forward and simply a case of mistakenly flipped indices. However, the second issue (*initialization issue*), which affects global alignments only, has a more substantial impact on alignment optimality and correctness. There exist several distinct formulations based on Gotoh’s original algorithm. Some of these are equivalent to Gotoh’s algorithm, while others require additional assumptions to yield correct results. For instance, Durbin describes an algorithm that, by design, only computes alignments where an insertion can not be directly followed by a deletion and vice versa [6]. The algorithm is correct, if some restrictions are imposed on the affine gap penalty and scoring matrix values. A sufficient condition is that the highest mismatch penalty is at most twice the gap extension penalty. Incidentally, on page 31 of [6], Durbin states this condition. On page 30 however, a different condition is given. For the latter, it is easy to show, that the condition is *not* sufficient for ensuring that insertions can not be followed by deletions in the optimal alignment.

All of the above generates confusion in the implementation of global alignment methods. Gotoh’s initialization error is present in standard textbooks (such as [9]) and in a plethora of online teaching material. Of the implementations we analyzed, some yield erroneous results, while others implicitly place additional assumptions on the alignment (e.g., no insertion can follow a deletion). This means that, the same two sequences can yield different alignments, depending on the software that is being used.

**Overview.** First, we give a description of Gotoh’s algorithm (Section 2.1), as it represents the cornerstone for constructing pairwise sequence alignments. Then, we present a detailed analysis of the errors that were introduced in the original paper and show how to avoid them (Section 2.2). Last, we assess the impact of these errors by listing books, implementations, and online lecture slides that either contain Gotoh’s mistake (books and lecture slides) or yield sub-optimal alignments (implementations). For lecture slides, we quantify the impact of the error, by the ratio of correct to incorrect presentations, and to lecture slides, where a formal initialization is missing altogether.

## 2 The Gotoh extension

To illustrate the two error types, we first recapitulate Gotoh’s algorithm for alignments with affine gap penalties. We use the same notation as in Gotoh’s original paper.

### 2.1 Gotoh’s algorithm

Let *w_k_* = *uk* + *v* (*u* ≥ 0, *v* ≥ 0) be the gap penalty for a gap of length *k*, where *v* is the *gap opening* penalty and *u* is the *gap extension* penalty. Let *A* = *a*_1_*a*_2_ *… a_M_* and *B* = *b*_1_*b*_2_ *… b_N_* be the two sequences we want to align. Further, assume that a weighting function *d*(*a_m_, b_n_*) is given to score an aligned pair of residues *a_m_* and *b_n_*. Typically, *d*(*a_m_,b_n_*) ≤ 0 if *a_m_* = *b_n_*, and *d*(*a_m_,b_n_*) > 0 if *a_m_* ≠ *b_n_*. The NW algorithm calculates the cells of a dynamic programming matrix *D_m,n_* using the recursion:

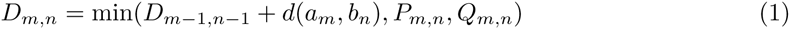

where

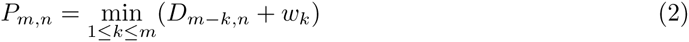

and

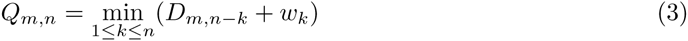

Here, *D_m,n_* is the score of a globally optimal alignment of the first *m* residues of *A* with the first *n* residues of *B*. *P_m_,_n_* is the score of an optimal alignment of the first *m* residues of *A* with the first *n* residues of *B* that ends with a deletion of at least one residue from *A*, such that *a_m_* is aligned with the gap symbol. Finally, *Q_m_,_n_* is the score of an optimal alignment of the first *m* residues of *A* with the first *n* residues of *B* that ends with an insertion of at least one residue from *B,* such that *b_n_* is aligned with the gap symbol. Although, at first sight, *P_m_,_n_* and *Q_m_,_n_* appear to require *m −* 1 (or *n −* 1) steps, they can be obtained in a single step via the following expansion of the recursive formulation:

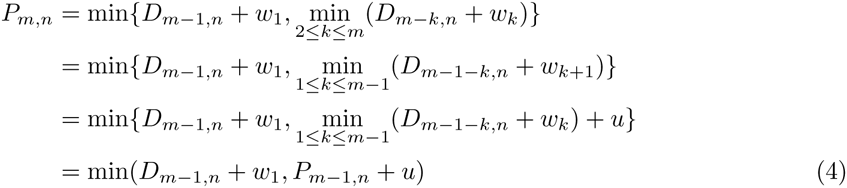

The same applies analogously to *Q_m,n_*:

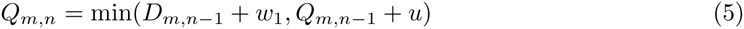

### 2.2 Mistakes in the Original Gotoh Algorithm

We have found two mistakes in the original Gotoh paper [8]. With respect to the initialization, Gotoh states:

“At the beginning of the induction, one may set *D_m_,*_0_ = *P_m_,*_0_ = *w_m_*(1 ≤ *m ≤ M*), and *D*_0_*,_n_* = *Q*_0_*_,n_* = *w_n_*(1 ≤ *n ≤ N*). Alternatively, *D_m_,*_0_ = *P_m_,*_0_ = 0 and *D*_0,_*_n_* = *Q*_0,_*_n_* = *w_n_*, or *D_m_,*_0_ = *P_m_,*_0_ = 0 and *D*_0_*,_n_* = *Q*_0_*,_n_* = 0 may be chosen in searching for the most locally similar subsequences …”.

Note that, the second sentence (at least the second part of it) refers to local alignments which are *not* affected by the error. Apart from the two errors we present in this section, there are additional issues in Gotoh’s paper, particularly in the description of the matrix traceback. In 1986, Altschul gave a detailed description of traceback issues introduced by Gotoh which can lead to sub-optimal alignments as well. For more information and examples see [1].

**Index Issue.** The first apparent mistake is that wrong indices are used for initializing the *P* and *Q* matrices. Initially, the entries *P_m_,*_0_ and *Q*_0_,*_n_*, as well as *D_m_*,_0_ and *D*_0_,*_n_* (for 1 ≤ *m ≤ M*, 1 *≤ n ≤ N*) are assigned some values. However, this is inconsistent with the recursions defined in equations 4 and 5. Consider computing the following entry *P*_1_,_1_ of *P* (or *Q*_1,1_ of *Q*). Equation 4 then reads as follows:

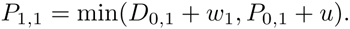

Here *D*_0,1_ is defined but *P*_0,1_ is *not* defined. However, *P*_1,0_ *is* defined, so this is a simple case of flipped indices. The same applies to matrix *Q*.

**Initialization Issue.** The more substantial problem are the actual values that are assigned to initialize *P* and *Q*. For global alignments, Gotoh proposes to initialize *D*_0_*_,n_* = *Q*_0_*_,n_* =*w_n_* and *D_m,_*_0_ = *P_m,_*_0_ = *w_m_* (for 1 ≤ *m ≤ M*, 1 ≤ *n ≤ N*). Correcting the indices for *P* and *Q* we obtain *D*_0_*_,n_* = *P*_0_*_,n_* =*w_n_* and *D_m,_*_0_ = *Q_m,_*_0_ = *w_m_*. The value *D*_0,0_ is defined as *D*_0,0_ = 0. Let us consider *P*_1_*,_i_* as defined in Equation 4 for some *i* ∈ [1, *N*]:

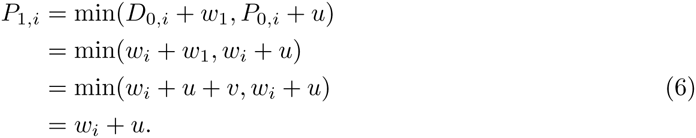

Similarly, for *j* ∈ [1, *M*]:

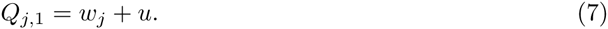

To illustrate why this result is wrong, we consider a simple one nucleotide example. Let *A* = *a*_1_ and *B* = *b*_1_. Further let *d*(*a*_1_*, b*_1_) := 5, the gap opening penalty *v* := 2, and the gap extension penalty *u* := 1. Now

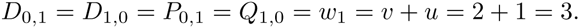

Thus, by equations 6 and 7 we obtain,

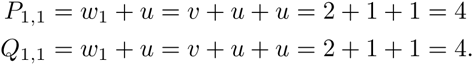

Plugging these values into Equation 1 we obtain

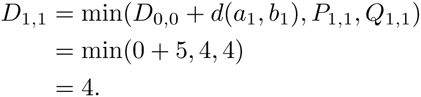

This implies that, the best alignment for *A* and *B* is:

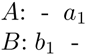

or

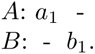

However, the actual correct score for both of these alignments is *w*_1_ + *w*_1_ = 3 + 3 = 6 ≠ 4. Aligning *A* and *B* as

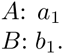

yields a score of *d*(*a*_1_, *b*_1_) = 5 < 6. Thus, conducting the initialization as proposed by Gotoh yields a sub-optimal solution for this simple example. Nonetheless, there is a straight-forward solution to this problem. We need to initialize the values for *P* and *Q* as *P*_0_*_,n_* ≥*w_n_* + *v* and *Q_m,_*_0_ ≥*w_n_* + *v* (for 1 ≤ *n* ≤ *N*, 1 ≤ *m* ≤ *M*) to obtain the correct, optimal alignment score. If *P*_0_*_,n_* =*w_n_* + *v* we can re-state Equation 6 as:

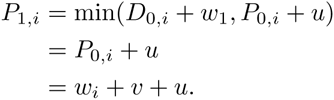

For *P*_0_*_,n_* > *w_n_* + *v* we get

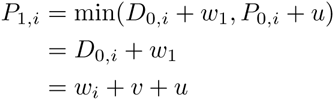

as well. A popular choice for *P*_0_*_,n_*, in publications by authors that seem to be aware of this issue, is *P*_0_*_,n_* := ∞ (see for example [1, 17]). A similar choice can be made for Q.

Using the corrected formula for our simple example of *A* = *a*_1_, *B* = *b*_1_, *d*(*a*_1_, *b*_1_) = 5, *v* = 2, and *u* = 1, we see that the values are correctly computed.

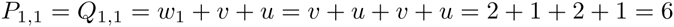

By Equation 1 we get

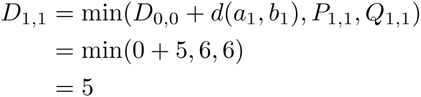

which is the correct result.

The values of *P*_1,_*_k_* (and analogously *Q_k_*,_1_) need to contain two gap opening penalties. By definition, they should represent the score of an optimal alignment of the first residue of *A* with the first *k* residues of *B* and end with a deletion of *a*_1_, that is, an alignment of *a*_1_ with the gap symbol. The resulting alignment will then always start with an insertion of the *k* first symbols of *B* followed by a deletion of the first symbol of *A*. However, according to Gotoh’s description, only a single gap opening penalty will be included.

## 3 Impact of the errors

Even though Gotoh’s paper was published over thirty years ago, the above error still persists in many papers and bioinformatics lectures. Furthermore, we are not aware of any previous work that specifically addresses the issues we have identified. Note that, there do exist publications that explain and/or implement a working or corrected version of the algorithm (e.g., [1, 5, 11, 14]). Other works either ignore this problem (e.g., [19]) or restrict values of *v, u,* and *d*(*a, b*) such that the issue disappears. For example, in 1972 Sankoff [15] originally solved the problem only for *u* = *v* = 0, and Durbin [6] gives an algorithm that performs well if 2*u* is greater than the highest value of *d*. Even though, some authors correct these mistakes on their own, numerous other publications, textbooks, and lecture notes still use the initial, incorrect, description. In the following, we list textbooks and lecture slides that contain the error. Further, we list software packages that yield sub-optimal alignments due to the issues described here or because of other conceptual errors. Note that, all open source software packages and implementations listed are available at http://www.exelixis-lab.org/web/software/alignment/.

### Books

The following two standard text books contain the initialization error.

– “Algorithms on Strings, Trees, and Sequences” by Gusfield, 2009 [9],
– “Introduction to Computational Biology” by Waterman, 1995 [22].

Fortunately, several books exist that contain a correct description of a global alignment algorithm, for instance [17].

### Software

**NW-align.** The alignment program **NW-align^4^** (e.g. discussed in [23]) shows the behavior described in Section 2.2 when aligning `GGTGTGA` with `TCGCGT.` **NW-align** assigns a score of −11 for gap opening and −1 for gap extension. Note that, the interpretation of affine gap costs is slightly different from Gotoh’s definition. Here, a gap of length k contributes a penalty of “−11 − (*k* − 1)” instead of “−11 − *k*” as defined in Section 2.1. **NW-align** produces the following alignment:

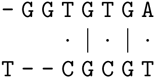

where the mismatch penalties are defined as *d*(*T, C*) := −1, *d*(*A, T*) := 0 and *d*(*G, T*) := −2. The score for the matches is defined as *d*(*G, G*) := 6. Thus, the score for this alignment is −11 − 11 −1 − 1 + 6 − 1 + 6 + 0 = −13. Considering the alignment

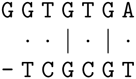

we can see that the result obtained by **NW-align** is sub-optimal, since the above alignment has a better score of −11 − 2 − 1 + 6 − 1 + 6 + 0 = −3.

**Bio**++. Bio++[7] is a **C++** library for Bioinformatics that includes methods for sequence comparison. The implementation of the Needleman-Wunsch-Gotoh method in the library can also generate suboptimal alignments. Aligning the sequences `AAAGGG` and TTAAAAGGGGTT by assigning 0 for a match, −1 for a mismatch, −5 for gap opening, and −1 for gap extension with the command

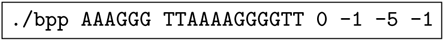

yields the following alignment with a score of −20:

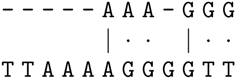

However, the following alignment has a better score of −15:

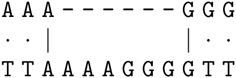

The sequences and parameters used here, are the same as used by Altschul [1] to demonstrate the error in Gotoh’s description of the traceback method. Interestingly, we observed another irregularity using Bio++. Running the implementation with the following options:

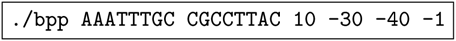

where the third argument (10) is the match score, the forth argument (−30) is the mismatch score and the last two arguments are the gap opening (−40) and extension costs (−1), yields the alignment.

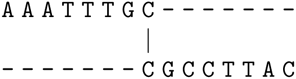

Surprisingly, flipping the input sequences

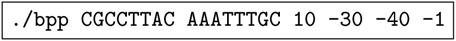

yields a different alignment with a different score:

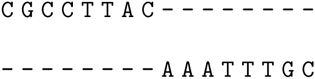

Nonetheless, both alignments are sub-optimal, since the alignment

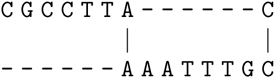

yields a better score of −72 (compared to −84 and −96 respectively).

**T-Coffee.** The **T-Coffee** package [13] for sequence alignment also implements the Gotoh algorithm. The command line used to produce the results below is

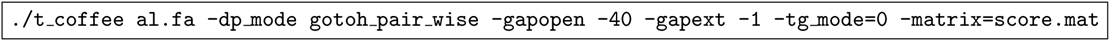

where `al.fa` contains the sequences TAAATTTGC and TCGCCTTAC. The gap opening penalty is −40, the gap extension penalty −1. The file `score.mat` defines a match score of 10 and a uniform mismatch score of −30. The resulting alignment as computed with T-Coffee is:

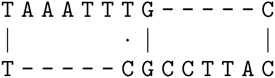

This alignment is sub-optimal. Consider the following alternative alignment:

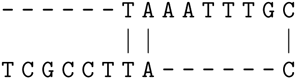

For the given parameters, the alignment returned by **T-Coffee** has a score of −90. However, the alternative alignment above, has a score of −62.

It might well be that the error in the pair-wise alignment also affects the multiple sequence alignment (MSA) algorithm in T-Coffee. However, T-Coffee does not only execute sequence-sequence, profile-sequence, or profile-profile alignments steps in the progressive MSA algorithm, but also uses additional concepts (e.g., the alignment information library). Therefore, it was not possible to reliably assess if this errors also affects the MSA procedure.

**FOGSAA.** The authors in [2] describe a branch-and-bound algorithm for global alignment that outperforms (in terms of speed) any optimal global alignment method including the widely used NW algorithm. Upon request via email, the authors provided us their implementation. To assess the correctness and speed of FOGSAA, the authors compared it to their own re-implementation of the NW algorithm. However, we obtained sub-optimal solutions when using this NW implementation to globally align sequences with affine gap penalties. For instance, given the sequences AAATTTGC and CGCCTTAC with the parameters *match* 10, *mismatch* −30, *gap opening* −40 and *gap extension* −1, we obtain the following alignment:

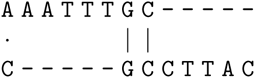

with a score of −100. The command we used is:

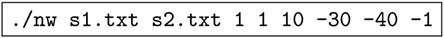

However, the following alignment is the optimal solution for this example:

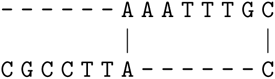

with a score of −72.

**HUSAR, MATLAB & BioPython.** Several implementations make the assumption that an insertion can not be followed directly by a deletion (or vice versa) in the optimal alignment. An algorithm that performs well (i.e., generates optimal alignments) under this assumption is the one by Durbin [6]. **HUSAR** is the information system of the DKFZ (German Cancer Research) and comprises several applications for sequence analysis. One such application is **GAP**, which performs pairwise sequence alignment and allows for affine gaps. While experimenting with it, we found that, **GAP** yields optimal alignments under the assumption that an insertion cannot follow a deletion (or vice versa). For instance, given a match score of 10, a mismatch of −30, gap opening −25, and gap extension −1, it generates the following alignment

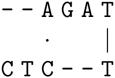

with score −74. The parameters are passed with:

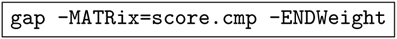

where `-MATRix` is the substitution matrix file name and `-ENDWeight` ensures that end gaps are also penalized. Assuming that, insertions and deletions can not reside immediately next to each other, this *is* the optimal solution. However, if we omit this assumption, the optimal alignment is

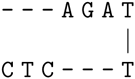

with a score of −46.

The corresponding function (*nwalign*()) in **MATLAB**^5^ yields an equivalent (in terms of alignment score) solution to **GAP**:

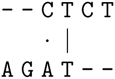

The **MATLAB** call is:

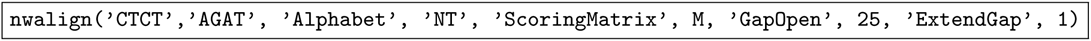

Note that, **MATLAB** returns a score of −72 for this alignment. This is due to the different possible interpretations of affine gap scores. That is, a gap of length *k* can contribute to the score with *v* + (*k* − 1)*u* instead of *v + ku*. Alternatively, one can apply a gap opening penalty of −26 to get the score of −74 reported by **GAP** for this alignment. The module **pairwise2**^6^ of the Biopython library [3] behaves analogously. The function

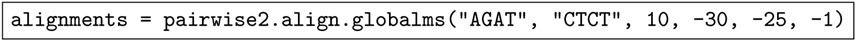

also yields alignments (including those found by **GAP** and **MATLAB**) with a score of −72. All three software packages do apparently not allow for insertions that are immediately followed by deletions. However, they do accept input values for which the optimal alignment does not exhibit this property.

**nwalign.** The **nwalign^7^** implementation is a python library (actually written in C) which implements global alignment with affine gaps. In some cases, it produces sub-optimal alignments as well. Again, consider the example of AGAT and CTCT. Given the same setup that we used for **HUSAR** (**GAP**), that is, a match score of 10, mismatch of −30, gap opening −25, and gap extension −1. The command:

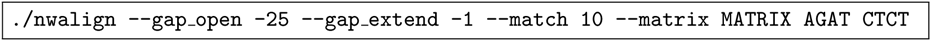

generates the correct alignment:

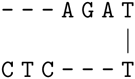

However, changing the scoring scheme to penalize opening a gap with −30 instead of −25 generates the following sub-optimal alignment:

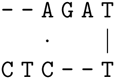

### Lecture slides

To further quantify the impact of the problem, we classified 31 lecture slides reported as the most popular results of Google search for the terms *global alignment, affine gaps, Needleman-Wunsch, Gotoh Algorithm,* into three distinct categories: *Correct, incomplete* and *wrong.* We observed that the majority (50%) of the slides (16 lectures, see Appendix A) are incomplete, since the initialization of the matrices is not explicitly given. Of course, lecture slides are only a part of the actual lectures. Hence, from the available resources we can not judge with certainty, whether an initialization (correct or incorrect) was presented to the students, for example orally, or via additional course material. Approximately 25% of the slides (7 lectures, see Appendix B) are correct. That is, a quadratic time algorithm is presented and a correct initialization is given. Slides that describe algorithms which make additional assumptions (e.g., Durbin [6]) are classified as correct if the initialization is correct for that particular case. Approximately 25% of the slides (8 lectures, see Appendix C) are wrong, that is, an incorrect initialization as described in Section 2.2 is provided. Other mistakes, such as stating incorrect conditions for avoiding subsequent insertions and deletions in the optimal alignment, are not counted as mistakes here. Slides that only describe the algorithm for locally aligning two sequences, without giving an algorithm for globally aligning sequences were discarded.

A List of incomplete lectures (16)
http://www.cs.utoronto.ca/~brudno/csc2427/Lec8Notes.pdf
ftp://statgen.ncsu.edu/pub/thorne/bioinf2/gotoh.pdf
http://www.cs.umd.edu/class/fall2011/cmsc858s/Gap_Scores.pdf
http://math.mit.edu/classes/18.417/Slides/alignment.pdf
http://users.ece.utexas.edu/~hvikalo/ee381v/lecture5h.pdf
http://ls11-www.cs.uni-dortmund.de/people/rahmann/teaching/ws2008–09/GrundlegendeBioinformatik/skript.pdf
http://www.csie.ntu.edu.tw/~kmchao/bioinformatics13spr/alignment.ppt
http://labs.bio.unc.edu/Vision/courses/162F02/03.pair.align.ppt
http://labs.bio.unc.edu/Vision/courses/162F02/04.mult.align.ppt
http://web.calstatela.edu/faculty/nwarter/courses/bioinfo/Bioinformatics_Sequence_Align_003.ppt
http://robotics.stanford.edu/~serafim/cs262/Slides/Lecture3.ppt
http://bioinfo.ict.ac.cn/~dbu/AlgorithmCourses/Lectures/Lec6-EditDistance.pdf
http://thor.info.uaic.ro/~ciortuz/SLIDES/pairAlign.pdf
http://www.cs.rice.edu/~nakhleh/COMP571/Slides/SequenceAlignment-PairwiseDP.pdf
http://www.cs.tau.ac.il/~bchor/CG09/CG2-alignment.ppt
http://www.cs.bilkent.edu.tr/~calkan/teaching/cs481/slides/cs481-Week4.2.pdf
http://angom.myweb.cs.uwindsor.ca/teaching/cs558/558-Lecture3.pptx

List of correct lectures (7)
http://ab.inf.uni-tuebingen.de/teaching/ws06/albi1/script/pairalign_script.pdf
http://www3.cs.stonybrook.edu/~rp/class/549f14/lectures/CSE549-Lec04.pdf
http://www.bioinf.uni-freiburg.de/Lehre/Courses/2014_SS/V_Bioinformatik_1/gap-penalty-gotoh.pdf
http://www.comp.nus.edu.sg/~ksung/algo_in_bioinfo/slides/Ch2_sequence_similarity.pdf
http://www.cs.cmu.edu/~ckingsf/class/02–714/Lec08-gaps.pdf
http://www.csie.ntu.edu.tw/~kmchao/seq11spr/Presentation_Sequence-final.pptx
http://wwwmayr.informatik.tu-muenchen.de/lehre/2009SS/cb/slides/CB1–2009–06–19.pdf

List of lectures containing mistake (8)
http://math.ucdenver.edu/~billups/courses/ma5610/lectures/lec4.pdf
http://users-cs.au.dk/cstorm/courses/AiBS_e14/slides/AffineGapcost.pdf
http://www.cise.ufl.edu/~cap5510fa13/02-CAP5510-Fall13.pptx
http://www.cs.uku.fi/~kilpelai/BSA05/lectures/print10.pdf
http://www.cse.msu.edu/~torng/Classes/Archives/cse960.01/Lectures/SequenceAlignment.ppt
http://www.haverford.edu/biology/GenomicsCourse/manduchi.ppt
http://www.site.uottawa.ca/~lucia/courses/5126–10/lecturenotes/03–05SequenceSimilarity.pdf
https://www.site.uottawa.ca/~turcotte/teaching/csi-5126/lectures/04/handouts.pdf

## Footnotes

4 Y. Zhang, http://zhanglab.ccmb.med.umich.edu/NW-align

5 ©2015 The Math Works, Inc. MATLAB and Simulink are registered trademarks of The Math Works, Inc. See www.mathworks.com/trademarks for a list of additional trademarks. Other product or brand names may be trademarks or registered trademarks of their respective holders.

6 Available at http://biopython.org/DIST/docs/api/Bio.pairwise2–module.html

7 This program (*nwalign*) is available at https://pypi.python.org/pypi/nwalign/

